# Integrating single-cell and bulk transcriptomic perturbation resources reveals complementary therapeutic spaces for drug repurposing

**DOI:** 10.64898/2026.07.17.739227

**Authors:** Enock Niyonkuru, Umair Khan, Xinyu Tang, Laura Almonte, Eden Chun, Brenda Ametepe, Carlota Pereda Serras, Boris Oskotsky, Brice Gaudillière, David K. Stevenson, Jessica Neely, Linda C Giudice, Tomiko Oskotsky, Marina Sirota

## Abstract

Transcriptome-based drug repurposing can accelerate therapeutic discovery, but is limited by fragmented resources, inconsistent quality control, and reliance on single perturbation databases. We developed CDRPipe (**C**omputational **D**rug **R**epurposing **Pipe**line), a unified framework that interrogates disease signatures against drug perturbation signatures generated by distinct experimental technologies. Specifically, CDRPipe harmonizes microarray perturbation profiles from the Connectivity Map (CMap; 1,968 quality-filtered experiments) with pseudo-bulk profiles derived from large-scale single-cell RNA sequencing experiments in the Tahoe-100M database (56,827 experiments). CDRPipe standardizes preprocessing, computes rank-based connectivity scores, and evaluates significance using empirical null models. We applied CDRPipe to 233 curated disease signatures from GEO and CREEDS and evaluated performance using known drug-disease associations from Open Targets. Single-cell-derived pseudo-bulk profiles recovered more annotated therapeutics than microarray profiles (median recall 50.0% vs. 6.2%; Wilcoxon p < 10⁻¹¹), thought these differences partly reflect differences in drug library composition and clinical annotation coverage. Importantly, the two resources were highly complementary, with only 3.5% overlap in recovered drugs, indicating that integrating predictions across independent perturbation resources expands therapeutic coverage and enables identification of high-confidence consensus candidates. Case studies in autoimmune disease and endometriosis further demonstrate that CDRPipe recovers clinically relevant therapies while revealing technology-dependent patterns of discovery. These results show that integrating heterogeneous transcriptomic perturbation resources improves the robustness and interpretability of transcriptional drug repurposing.

**One Sentence Summary:** Integrating drug perturbation resources from distinct transcriptomic platforms improves the robustness and accuracy of drug repurposing predictions.

## INTRODUCTION

The contemporary landscape of pharmaceutical research and development (R&D) is characterized by a paradoxical trend often referred to as “Eroom’s Law”, the observation that drug discovery is becoming slower and more expensive over time, despite improvements in technology (*1*). Developing a therapeutic agent de novo is a capital-intensive endeavor, estimated to cost between $2 billion and $3 billion and requiring 10 to 17 years to move from target identification to market approval (*2*). Compounding this financial burden is a high attrition rate; approximately 90% of drug candidates entering clinical trials fail to achieve regulatory approval, often due to lack of efficacy or unforeseen toxicity in late-stage development (*3*). This systemic inefficiency leaves a vast array of diseases, particularly rare conditions and complex chronic disorders, without effective targeted treatments.

In response to these challenges, drug repurposing, the identification of novel therapeutic indications for existing drugs with established safety profiles, has emerged as a critical strategy to accelerate therapeutic discovery and reduce translational risk (*4, 5*). Historically, repurposing successes were largely unanticipated, driven by clinical observation of off-target effects, as exemplified by the redeployment of sildenafil for erectile dysfunction or thalidomide for multiple myeloma (*1, 6*). However, the explosion of high-throughput “omics” data has catalyzed a paradigm shift toward systematic, computational drug repurposing (*5, 7*, *8*).

Central to this modern approach is the hypothesis of transcriptional reversal (or signature reversion). This principle posits that if a disease state is characterized by a specific gene expression signature (a set of up- and downregulated genes), a small molecule capable of inducing the opposite transcriptional profile may neutralize the disease phenotype and restore a healthy physiological state (9). This concept was pioneered by the Connectivity Map (CMap) project, which provided the first large-scale reference database of drug-induced transcriptional perturbations, allowing researchers, including our own team, to systematically connect small molecules, genes, and diseases (*10–12*).

Despite the conceptual elegance and early successes of transcriptomic drug repurposing, the field faces persistent limitations related to data heterogeneity, platform bias, and biological resolution. In bulk transcriptomic experiments, gene expression is measured across all cells present in a tissue or sample and represented as a single composite profile. As a consequence, transcriptional programs originating from rare but disease-driving cell populations can be masked by signals from more abundant but less relevant cell types, reducing the sensitivity of bulk-derived signatures to capture true disease mechanisms. This dilution of cell-type-specific signals can cause therapeutically relevant compounds, those targeting the actual disease-driving pathways, to fail to show significant transcriptional reversal against the composite bulk signature, leading to missed therapeutic associations. At the same time, compounds that induce broad transcriptional suppression or cytostatic responses may be artifactually favored over those with more targeted mechanisms of action (*13*). Furthermore, reliance on a single perturbation technology can introduce platform-specific biases that limit generalizability (*14*). The emergence of large-scale single-cell RNA sequencing (scRNA-seq) perturbation databases presents an opportunity to diversify the transcriptional resources available for drug repurposing (*14–16*). While single-cell perturbation profiling does not directly resolve cell-type heterogeneity in bulk disease signatures, it offers complementary advantages: broader genomic coverage, a wider diversity of cellular contexts for measuring drug responses, and the ability to capture drug-induced transcriptional effects that bulk microarray platforms may fail to detect. For example, Tahoe-100M is a massive single-cell perturbation atlas comprising approximately 100 million scRNA-seq profiles that capture transcriptional responses to small-molecule treatments across dozens of cancer cell lines. Yet, the integration of these high-dimensional, disparate data types remains a significant computational bottleneck (*17*), often forcing researchers to choose between the extensive drug coverage of legacy bulk databases and the granular resolution of modern single-cell platforms.

To address these challenges, we developed CDRPipe, a unified framework for transcriptional drug repurposing that enables direct comparison and integration of transcriptional perturbation signatures derived from heterogeneous measurement technologies. CDRPipe standardizes preprocessing, applies consistent nonparametric connectivity scoring, and evaluates statistical significance within a shared empirical framework. We hypothesized that therapeutically relevant compounds would exhibit consistent transcriptional reversal across independent perturbation resources, and that pseudo-bulk signatures derived from single-cell experiments would improve the recovery of known drug-disease associations compared to microarray-based approaches alone.

By applying CDRPipe to 233 curated disease signatures from CREEDS (Crowd-Extracted Expression of Differential Signatures) (*18*) and validating predictions against curated drug-disease knowledge bases, such as Open Targets (*19*), we demonstrate that multi-database integration substantially enhances the robustness, interpretability, and translational relevance of transcriptional drug repurposing. We further illustrate the utility of this framework through case studies in autoimmune diseases, where we recover established immunomodulators and identify novel targeted therapies, and in endometriosis, where we replicate prior findings while uncovering new enzyme-centric therapeutic candidates.

## RESULTS

### Study Overview

To systematically evaluate the impact of cellular resolution on drug repurposing, we integrated heterogeneous transcriptional resources into a unified analytical framework (***Fig. 1***). On the disease side, we utilized disease transcriptional signatures obtained from the CREEDS resource (*18*), comprising manually curated differential gene expression profiles derived from public studies in the Gene Expression Omnibus (GEO) (*20*). Each signature contrasts diseased tissues or cells with matched normal or control samples within the same study. Signatures were generated through a large-scale crowdsourcing effort with manual selection of samples, standardized disease annotation, batch effect correction using surrogate variable analysis, and differential expression prioritization via the Characteristic Direction method. All signatures were derived from microarray-based transcriptomic experiments (*18*). We analyzed 233 manually curated disease signatures, each defined by sets of upregulated and downregulated genes, with a mean size of approximately 975 genes (range: 38-4,867) prior to filtering and 419 (range: 2 - 2,525) after applying significance and fold-change thresholds (***Fig. 1***). For therapeutic profiles, we employed two distinct perturbation databases: the Connectivity Map (CMap)(*12*) and Tahoe-100M (***Table 1***) (*16*). CMap provides direct microarray expression profiles of drug-treated cells; after quality-control filtering based on inter-replicate consistency (Pearson correlation r ≥ 0.15 between replicate signatures for the same drug-dose-cell line combination), we retained 1,968 high-quality drug perturbation experiments out of 6,100 (32.3%), spanning 13,071 unique gene features across five cell lines and 1,309 unique drugs (***Fig. 1***). Tahoe-100M is a single-cell perturbation atlas (scRNA-seq based), from which we used 56,827 drug-response experiments aggregated into pseudo-bulk expression signatures; no experiment-level filtering was required for Tahoe-100M (given its inherently high data quality), and the resulting profiles covered 22,168 unique genes. Tahoe-100M covers 50 cell lines and with 379 unique drugs. Tahoe-100M and CMap have 85 drugs and 12,544 genes in common, but no cell lines are shared (***Fig. 1F*, *Table 1***).

**Figure 1:**
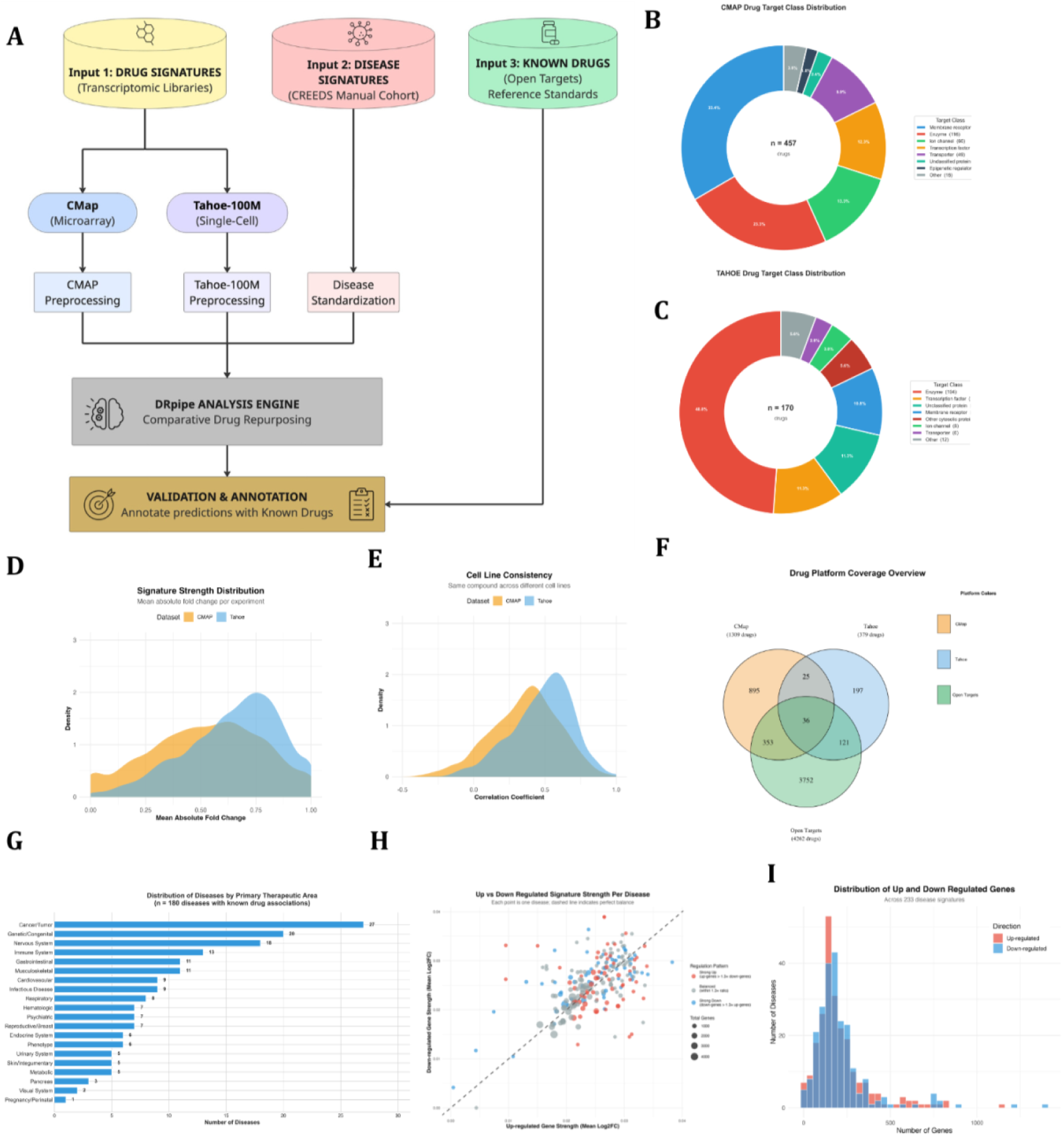
Overview and data descriptors. **Study Workflow, Dataset Characteristics, and Quality Control Metrics.(A)** The CDRpipe framework integrates transcriptomic drug perturbation libraries from **CMap** (microarray-based) and **Tahoe-100M** (single-cell RNA-seq) with disease signatures from the **CREEDS** manual cohort. Following platform-specific preprocessing and quality control (QC), the Comparative Drug Repurposing (CDR) engine predicts therapeutic associations, which are subsequently validated against known indications from **Open Targets**. **(B)** Target class distribution for 457 CMap drugs, showing a receptor-oriented profile dominated by membrane receptors (33.4%). **(C)** Target class distribution for 170 Tahoe-100M drugs, revealing an enzyme-centric bias (48.8%) reflecting the library’s focus on intracellular signaling. **(D)** Comparative signature strength distribution; Tahoe-100M exhibits a shift toward higher fold changes compared to CMap. **(E)** Cross-cell line consistency, showing comparable correlation coefficients for compounds profiled across varying lineages. **(F)** Venn diagram of the chemical space overlap. Only 36 drugs are shared across all three datasets, underscoring the complementary pharmacological coverage provided by the multi-platform approach. **G)** Distribution of the 180 diseases analyzed, categorized by primary therapeutic area. Oncology (n=27) and Genetic/Congenital disorders (n=20) represent the largest cohorts**. (H)** Correlation between up-regulated and down-regulated gene strength. Most diseases cluster along the diagonal, indicating balanced regulatory signatures. **(I)** Distribution of gene counts per signature, confirming a balanced representation of up-regulated and down-regulated genes (mean = 209 each).

**Table 1:** Drug Signatures Comparison of Tahoe-100M and CMap Table. **Comparison of Drug Signature Characteristics between CMap and Tahoe-100M.** This table summarizes the key structural and experimental differences between the two perturbation resources integrated into the pipeline. It contrasts the scale of each dataset (number of unique drugs, genes, and expression profiles), the biological resolution (number of cell lines), and the underlying measurement technology (Microarray vs. Single-cell RNA-seq). The “Shared” column highlights the overlap between the libraries, specifically the 85 drugs and 12,544 genes common to both platforms, which serve as the basis for the comparative analysis.

| Category | CMap | Tahoe-100M | Shared |
| --- | --- | --- | --- |
| Unique drugs | 1,309 | 379 | 85 |
| Unique genes | 13,071 | 25,084 | 12,544 |
| Cell lines | 5 | 50 | 0 |
| Gene matrix dimensions | $\sim 6,100 \times 13,071$ | $56,827 \times 62,710$ | — |
| Technology | Microarray HG U133A | Single-cell RNA seq with aggregated pseudo bulk matrices |  |
| Total profiles | 6,100 experiment profiles | > 100,000,000 single cell profiles and 56,827 aggregated experiment profiles |  |
| Expression values per experiment | ranked probe set signatures | basemean, log2FC, p value, p adjusted |  |

CDRPipe integrates these heterogeneous resources within a unified analytical pipeline, applying uniform preprocessing and a nonparametric connectivity scoring method to quantify how strongly each drug’s transcriptomic profile reverses a given disease signature, with statistical significance assessed by empirical permutation-based null models. We applied this pipeline to all 233 disease signatures to generate drug repurposing predictions, and then validated the results using Open Targets, an external knowledge base of known disease-drug associations (Open Targets Platform, 2024 release) (*19*). Disease terms from CREEDS were matched to Open Targets indications via exact name matching (151 diseases, 64.8%) or synonym matching (52 diseases, 22.3%), yielding 203 diseases with at least one known therapy for validation (***Fig. 1F***). Altogether, Open Targets provided a benchmark set of thousands of known disease-drug links for these conditions (over 118,000 associations in total), of which 2,668 disease-drug pairs involved drugs present in the CMap/Tahoe-100M datasets and were evaluated in our analysis. For Open Targets, we include disease-drug pairs with evidence of development activity, including drugs in clinical trials across all phases. In summary, CDRPipe unifies disparate disease signatures and drug perturbation profiles (bulk and single-cell) into a single framework for systematic drug repurposing, and its performance is rigorously assessed against a large compendium of established disease-drug relationships.

### Prediction Recovery and Validation

We assessed the performance of our approach by calculating precision (proportion of predictions confirmed in Open Targets) and recall (proportion of recoverable known relationships successfully predicted). We note that Open Targets captures drugs with any recorded development activity, including early-phase clinical trial candidates, and is biased toward compounds with established clinical annotation; these metrics therefore reflect recovery of documented associations within the benchmark rather than absolute predictive accuracy. To account for overlapping annotations, diseases sharing identical therapeutic area combinations in Open Targets were grouped, with disease therapeutic areas referring to the broad clinical categories the disease falls under as defined by Open Targets. Tahoe-100M recovered more annotated drug-disease pairs than CMap (2,198 vs. 948), a 2.3-fold difference. Tahoe-100M also covered more diseases (171 vs. 155) and achieved a higher overall recovery rate, defined as the proportion of recoverable known drug-disease pairs (i.e., those where the drug exists in the platform’s library) that were successfully predicted with a significant reversal score, of 22.1% versus 18.1%. It is important to note that these differences partly reflect the composition of each drug library: Tahoe-100M contains many newer, clinically oriented small-molecule compounds that may have inherently higher overlap with Open Targets annotations, whereas CMap’s library includes older and more exploratory chemical entities.

We further quantified performance metrics for each disease, defining *I* as unique drugs predicted, *S* as predicted drugs validated in Open Targets, and *P* as the recoverable ceiling (known drugs present in the platform’s library). Precision was calculated as *S*/*I* and recall as *S*/*P*. Using lung adenocarcinoma as a representative example, Tahoe-100M predicted 61 unique drugs (*I* = 61), 10 of which were validated (*S* = 10), yielding a precision of 16.4%. Of the 48 known drugs for this condition, defined here as drugs with any recorded development activity in Open Targets, 12 existed in the library (*P* = 12); thus, the pipeline recovered 10 of 12 recoverable drugs, achieving 83.3% recall.

At the disease level across all 233 evaluated diseases, Tahoe-100M recovered a higher proportion of Open Targets-annotated drug-disease pairs than CMap (**Fig. 2**). Mean precision was 1.8% (SD 3.3%) for Tahoe-100M versus 1.3% (SD 2.8%) for CMap (measured across n=223 and n=208 diseases with at least one prediction, respectively) (**Fig. 2 A**). Among diseases with recoverable drugs (*P* > 0), Tahoe-100M achieved a mean recall of 47.3% (SD 38.5%) compared to 18.5% (SD 25.1%) for CMap (n=156 and n=165 diseases, respectively; **Fig. 2B**). These differences should be interpreted cautiously: they partly reflect the greater overlap between Tahoe-100M’s drug library and Open Targets annotations rather than representing a direct measure of predictive superiority. Nonetheless, many diseases achieved >20% recall with Tahoe-100M compared to substantially fewer with CMap, and multiple diseases achieved >50% recall with Tahoe-100M, a threshold not reached by any disease in CMap analyses (Fig. 2 C). Variable sample sizes reflect platform-specific prediction coverage and the filtering requirements for each metric (*I* > 0 for precision calculations and *P* > 0 for recall calculations).

**Figure 2:**
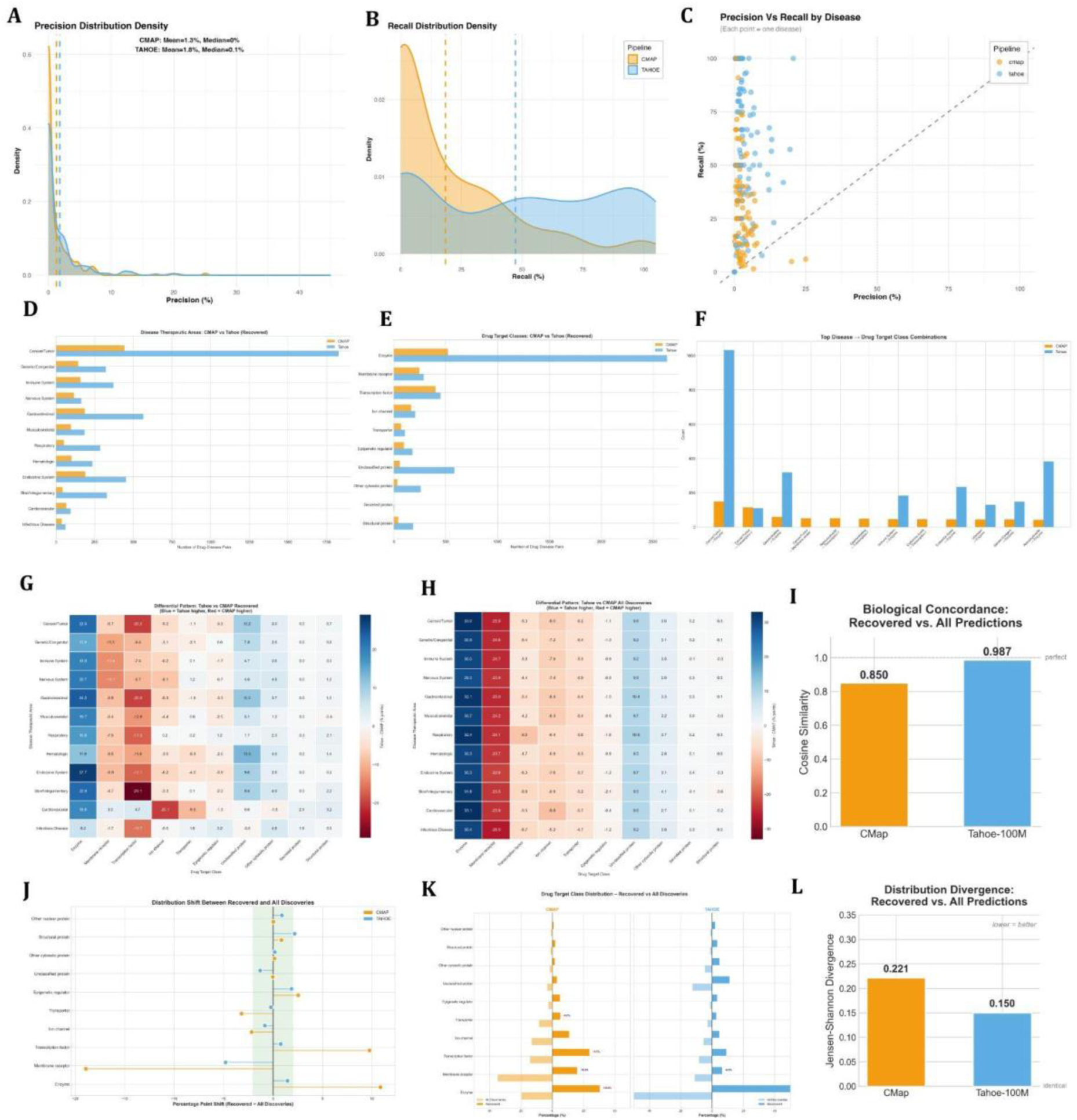
Prediction Results precision and recall, disease specific summaries and biological concordance. **Comparative Performance Benchmarking and Pharmacological Discovery Patterns. (A–C) Precision and Recall Performance Across Diseases**. Benchmarking of the CDRPipe engine using Open Targets as a gold standard.**(A) Precision distribution density** displaying right-skewed profiles for both platforms. Low mean precision (CMap: 1.3%; Tahoe-100M: 1.8%) reflects the inherent challenge of predicting validated relationships from transcriptomic signatures alone.**(B) Recall distribution density** for CMap (n=165 diseases) and Tahoe-100M (n=156). Dashed vertical lines indicate mean recall (CMap: 18.5%, Tahoe-100M: 47.3%), illustrating Tahoe-100M’s significantly superior recovery of known drug-disease relationships. (**C) Precision versus Recall scatter plot** by disease. Tahoe-100M (blue) clusters at higher recall values than CMap (orange) while maintaining comparable precision, demonstrating superior overall recovery of therapeutically validated drug-disease relationships.**(D–F) Platform-Specific Prediction Profiles**.**(D) Recovery by therapeutic area.** Tahoe-100M demonstrates superior recovery volume across most categories, with particularly strong enrichment in Oncology, Gastrointestinal, and Immune System disorders.**(E) Recovery by drug target class.** Tahoe-100M displays a distinct mechanistic bias toward Enzymes, reflecting its sensitivity to signaling pathway inhibitors, whereas CMap shows a more balanced distribution across Membrane Receptors and Ion Channels. **(F) Top Disease-to-Target combinations.** The “Cancer/Tumor → Enzyme” axis dominates Tahoe-100M’s performance, driven by the successful recovery of kinase inhibitors. In contrast, CMap recoveries are more evenly distributed across diverse mechanisms. **(G-L) Biological Concordance Between Validated and Novel Predictions**. **(G - H) Differential discovery patterns.** Heatmaps display the discovery frequency difference between Tahoe-100M (blue) and CMap (red) for **(G)** recovered (validated) and **(H)** all (total) computational predictions. The high similarity between panels indicates that Tahoe-100M’s novel predictions faithfully reflect the biological mechanisms of validated drugs. **(I) Cosine similarity** between recovered and all prediction target-class distributions for each platform. Tahoe-100M (0.987) approaches the theoretical maximum of 1.0, indicating near-identical mechanistic profiles, whereas CMap (0.850) shows greater divergence. **(J) Target class distributions** (Butterfly Chart). Comparison of “All Discoveries” (light) versus “Recovered” (dark) predictions. CMap shows notable shifts between discovery and validation phases, while Tahoe-100M distributions remain consistent. **(K) Distribution shift** (Lollipop Chart). Quantifies the percentage point shift between total discovery and validated recovery. CMap exhibits large deviations (e.g., -18.9% for membrane receptors), whereas Tahoe-100M clusters tightly within a ±2% “Minimal Shift” zone, demonstrating superior biological stability and predictive reliability. **(L) Jensen-Shannon divergence** between the same distributions. Lower values indicate greater similarity; Tahoe-100M (0.150) exhibits less distributional shift than CMap (0.221), confirming that Tahoe-100M’s novel predictions more faithfully recapitulate the target-class composition of its validated hits.

### Complementarity of Platforms and Mechanistic Profiles

The central finding of our cross-platform analysis is the striking complementarity between CMap and Tahoe-100M. Analysis of the overlap between drug-disease pairs predicted by each platform revealed low concordance (***Fig. 2D***). Among drug-disease pairs supported by Open Targets, only 144 were identified by both CMap and Tahoe-100M (Jaccard index 4.8%). When considering all predicted pairs, including those not supported by Open Targets, the overlap remained limited (173 pairs; Jaccard index 1.2%). Despite both platforms being applied to the same set of 233 disease signatures, the majority of predicted pairs were unique to a single platform, with each exhibiting different recovery volumes and enrichment patterns across therapeutic areas and drug target classes (***Fig. 2D, 2E***). This divergence is consistent with the limited chemical space overlap between the two drug libraries (***Fig. 1F***) and is further reflected in the distinct disease-to-target combinations favored by each platform (***Fig. 2F***). Together, these results indicate that CMap and Tahoe-100M sample largely non-overlapping therapeutic spaces rather than redundantly identifying the same candidates, and that the primary value of integrating both platforms lies in this complementarity.

While both platforms validated predictions across diverse therapeutic areas, with cancer/tumor being the largest category for both, Tahoe-100M’s recovered predictions were significantly enriched for oncology, with 28.7% of Tahoe-100M’s recovered drug-disease pairs falling in oncology compared to 14.3% for CMap (***Fig. 2D–2F***), consistent with its enzyme-centric drug library bias (***Fig. 1C***).

### Biological Concordance of Novel Predictions

To determine if novel predictions adhered to the same therapeutic logic as clinically validated drugs, we evaluated the biological concordance between “recovered” (validated) predictions and “all” discoveries by comparing their joint distributions across disease categories and drug target classes. Heatmaps of these distributions revealed that Tahoe-100M’s differential discovery patterns were highly similar between recovered and all predictions (***Fig. 2G, 2H***), yielding a cosine similarity of 0.987 and a Jensen-Shannon divergence of 0.150 (***Fig. 2I, 2L***). This consistency indicates that Tahoe-100M’s novel candidates closely mirror the mechanistic profile of its validated successes.

CMap showed lower concordance between its recovered and all discovery distributions (cosine similarity = 0.850; Jensen-Shannon divergence = 0.221; ***Fig. 2I, 2L***). The butterfly chart of target class distributions (***Fig. 2J***) shows that CMap exhibits notable shifts between the “all discoveries” and “recovered” profiles, particularly for membrane receptors, whereas Tahoe-100M distributions remain consistent. This is quantified in the lollipop chart (***Fig. 2K***), where CMap exhibits large percentage-point deviations between total discovery and validated recovery (e.g., −18.9% for membrane receptors), while Tahoe-100M clusters tightly within a ±2% shift zone. This higher divergence for CMap implies that it explores a broader chemical space, potentially identifying mechanistically novel relationships that differ from established clinical precedents. We note that these platform-specific mechanistic profiles partly reflect the inherent composition of each drug library (***Fig. 1B, 1C***): Tahoe-100M’s enzyme-enriched library naturally yields enzyme-centric predictions, while CMap’s receptor-heavy library favors receptor-targeted discoveries.

### Case Study 1: Autoimmune Diseases

To evaluate pipeline performance, we analyzed 18 autoimmune diseases mapped to Open Targets via exact name or synonym matching, details about the diseases available in Supplemental Table 1 (***Supplemental Table 1***). Note that the “known drugs” column in Supplemental Table 1 reflects all drugs with any recorded development activity for that indication in Open Targets (including clinical trial candidates at all phases), rather than only approved therapies; this accounts for the high counts observed for some diseases. Autoimmune diseases constitute a rigorous, biologically grounded benchmark for transcriptional drug repurposing, as their pathogenesis is driven by aberrant immune activation that manifests as reproducible gene expression signatures (*21*). Despite clinical heterogeneity, these disorders converge on shared inflammatory axes, including interferon signaling, cytokine-mediated activation, and T-cell dysregulation, allowing for a systematic evaluation of the pipeline’s ability to recover established immunomodulators (*22*). Furthermore, the persistence of treatment resistance in autoimmune conditions underscores the clinical urgency for scalable approaches that nominate candidates based on molecular reversal rather than phenotype alone. As shown in ***Fig. 3B***, Tahoe-100M significantly outperformed CMap in recovering known therapeutics in the Open Targets database (mean recovery rate: 76.5% vs. 20.9%; Wilcoxon signed-rank test p < 0.001; Cohen’s d = 2.22), representing a 3.7-fold improvement (***Fig. 3***). Disease-specific analysis revealed Tahoe-100M’s particular strength in inflammatory conditions: Crohn’s disease (75.0% vs 7.4%), psoriasis (100.0% vs 75.0%), and rheumatoid arthritis (64.0% vs 17.8%) (***Fig. 3A***). However, CMap demonstrated better performance for type 1 diabetes mellitus, which may reflect the fact that its treatment paradigm centers on glucose regulation rather than immunosuppression, potentially favoring CMap’s drug library composition.

**Figure 3.**
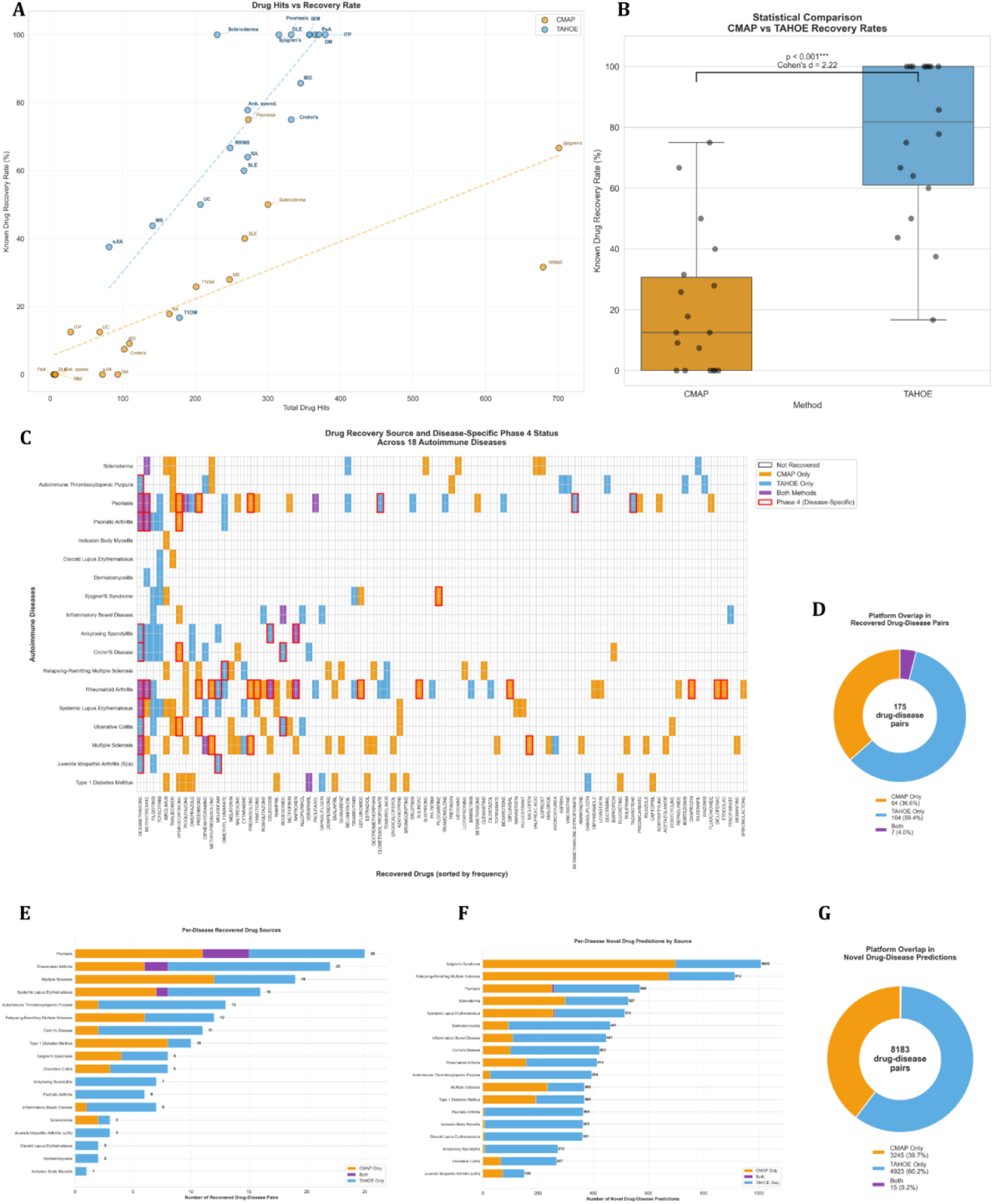
Case Study 1: Autoimmune Diseases. **Case Study: Autoimmune Diseases. (A) Drug Hits vs. Recovery Rate.** Scatter plot comparing the total number of predicted drug hits against the recovery rate of known therapeutics for 18 autoimmune diseases. Tahoe-100M (blue) demonstrates a strong positive correlation where increased predictions translate to higher recovery, whereas CMap (orange) shows higher variability and lower ceilings for recovery. **(B) Statistical Comparison.** Box plots quantifying the difference in known drug recovery rates. Tahoe-100M significantly outperforms CMap (Mean: 76.5% vs 20.9%; Wilcoxon signed-rank test p < 0.001, Cohen’s d = 2.22), representing a 3.7-fold improvement in identifying validated treatments. **(C) Drug Recovery Matrix and Clinical Validation.** Heatmap displaying the 94 unique drugs recovered across the autoimmune cohort. Cells are colored by platform source: **Orange** (CMap Only), **Blue** (Tahoe Only), and **Purple** (Both). **Red borders** denote drugs validated in Phase 4 clinical trials specifically for that disease indication. The matrix highlights the low overlap between platforms (sparse purple cells, ∼4.0%) and the high clinical relevance of the predictions, particularly for Tahoe-100M in diseases like Rheumatoid Arthritis and Psoriasis. **(D) Platform Complementarity (Recovered).** Donut chart showing the distribution of 175 recovered drug-disease pairs by platform source: Tahoe-only (104, 59.4%), CMap-only (64, 36.6%), and Both (7, 4.0%). **(E) Per-Disease Recovered Drug Sources.** Horizontal stacked bar chart displaying the per-disease breakdown of recovered drugs by platform source, sorted by total count. Psoriasis and rheumatoid arthritis dominate both in volume and in consensus predictions (purple), while several diseases (e.g., ITP, Crohn’s, ankylosing spondylitis) show exclusively single-platform recovery, underscoring the orthogonal therapeutic spaces sampled by each platform. **(F) Per-Disease Novel Drug Predictions.** Horizontal stacked bar chart showing the per-disease breakdown of novel (unvalidated) drug predictions by platform source. Sjögren’s syndrome (1,008) and relapsing-remitting multiple sclerosis (913) generate the most novel predictions, reflecting the large candidate pools available for these conditions. **(G) Platform Overlap in Novel Predictions.** Donut chart showing the distribution of 8,183 novel drug-disease pairs: CMap-only (3,245, 39.7%), Tahoe-only (4,923, 60.2%), and Both (15, 0.2%). The near-zero overlap (0.2%) in novel predictions is even more pronounced than in recovered drugs (4.0%), reinforcing that the two platforms sample largely non-overlapping therapeutic spaces.

We further validated these findings by examining disease-specific Phase 4 clinical trial data (***Fig. 3C***). Of the recovered drug-disease pairs, 26.3% (46 of 175) involved drugs validated in Phase 4 specifically for that indication. The most frequently recovered agents were cornerstone immunosuppressants, including dexamethasone (identified in 12 diseases), methotrexate (11 diseases), and hydrocortisone (7 diseases). Several robust disease-specific patterns emerged: rheumatoid arthritis demonstrated the highest concordance with Phase 4 data, recovering 15 distinct Phase 4-approved drugs. This is consistent with rheumatoid arthritis having one of the broadest therapeutic arsenals among autoimmune diseases, providing a larger validation set. Both platforms consistently predicted methotrexate, celecoxib, naproxen, dexamethasone, and meloxicam, representing greater therapeutic validation than other autoimmune conditions evaluated. Similarly, the psoriasis spectrum (including psoriatic arthritis) showed consistent recovery of dimethyl fumarate, tazarotene, and clobetasol propionate. Gastrointestinal autoimmune diseases (Crohn’s disease and ulcerative colitis) preferentially recovered locally-acting corticosteroids such as budesonide. Sjögren’s syndrome recovered pilocarpine and leflunomide, reflecting its distinct pathophysiology centered on exocrine gland dysfunction.

Despite Tahoe-100M’s overall superiority, the two methods showed high complementarity with only 4.0% overlap in recovered drug-disease pairs. Across 175 recovered drug-disease pairs, Tahoe-100M uniquely contributed 104 (59.4%), while CMap uniquely identified 64 (36.6%) that Tahoe-100M missed. Beyond validated associations, both platforms generated substantial novel predictions: CMap produced 4,900 novel (unvalidated) drug-disease pairs involving 198 drugs that had no Open Targets confirmation, while Tahoe-100M produced 8,766 novel pairs involving 32 exclusively novel drugs, a difference reflecting Tahoe-100M’s smaller but more clinically oriented library, where most drugs already have some Open Targets annotation (***Fig. 3F, 3G***). Drugs predicted by both methods showed 2.6-fold higher precision (5.2%) compared to single-method predictions (1.8 - 2.0%), suggesting consensus predictions represent higher-confidence repurposing candidates. Across the full panel of 18 autoimmune diseases, we identified 94 unique recovered drugs representing the combined outputs of both databases. Only 7 of 175 drug-disease pairs (4.0%) were identified by both platforms, reinforcing that distinct signals are captured by CMap and Tahoe-100M (***Fig. 3C, 3D***).

Qualitative analysis of the recovered candidates revealed distinct platform-specific therapeutic spaces. CMap excelled at recovering established medications, including NSAIDs, corticosteroids, and traditional immunosuppressants. In contrast, Tahoe-100M uniquely identified emerging targeted therapies, including JAK inhibitors (tofacitinib, filgotinib) across four diseases, BTK inhibitors (tirabrutinib), and SGLT2 inhibitors for diabetes. This difference partly reflects the composition of each drug library: Tahoe-100M includes newer, clinically oriented compounds not present in CMap, while CMap’s library is enriched for older, well-established pharmacological agents. These findings underscore the value of a multi-platform approach: each platform captures a distinct pharmacological space, and their integration maximizes overall therapeutic coverage. While the volume of novel candidates is substantial, all predictions are rank-ordered by their connectivity score, providing a principled prioritization for downstream evaluation. The high candidate counts partly reflect the use of uniform statistical thresholds across all 18 diseases; disease-specific parameter tuning could narrow the output but risks introducing subjective bias. Importantly, the ranking ensures that the most promising candidates, those with the strongest expression-reversal signatures, appear at the top, enabling systematic experimental follow-up in order of predicted efficacy rather than requiring exhaustive evaluation of the full list.

### Case Study 2: Endometriosis

To further validate our pipeline, we applied Tahoe-100M to endometriosis, benchmarking our results against the prior CMap-based analysis by Oskotsky et al (*6*). Endometriosis presents a unique opportunity for computational discovery due to the critical unmet need for non-hormonal therapies and the historical stagnation of traditional research and development (*23*). Biologically, however, it is an ideal candidate for this approach as the disease shares significant molecular hallmarks with malignancy, including invasion, angiogenesis, and metastasis (*24*). This transcriptional overlap is particularly advantageous given that the reference data in both Tahoe-100M and CMap are derived largely from cancer cell lines. We began by replicating the findings of Oskotsky et al (*6*) using the CDRPipe framework. We achieved 100% replication of their reported results using the original study parameters: a p-value less than 0.05, an absolute log2 fold change greater than 1.1, a random seed of 2009, and 1,000 permutations (***Fig. 4A***). Following this validation, we applied the same rigorous parameters to the Tahoe-100M pipeline (***Fig. 4B***). While Oskotsky et al. stratified signatures by disease stage and menstrual phase, we focused our comparative ranking on the “Unstratified” signature to identify robust, broad-spectrum candidates.

**Figure 4.**
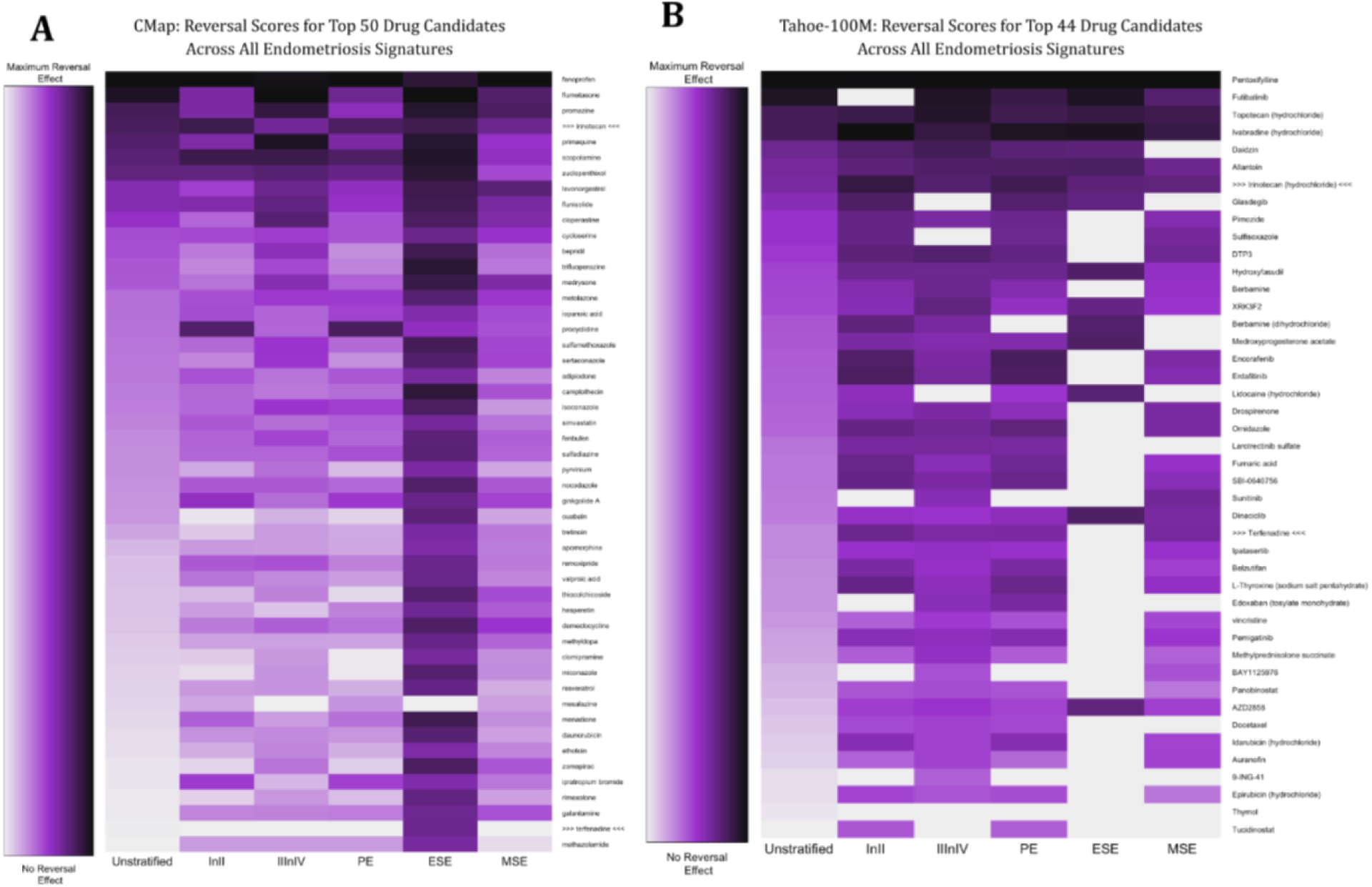
Case Study 2 : Endometriosis. **Comparison of Top Drug Repurposing Candidates for Endometriosis via CMap and Tahoe-100M Pipelines. (A) CMap Top 50 Candidates.** Heatmap displaying the top-ranked drugs identified using the CMap microarray database. Rows represent individual drugs, and columns correspond to the six endometriosis signatures (Unstratified, Stage I/II, Stage III/IV, Proliferative Phase, Early Secretory Phase, Mid-Secretory Phase). Darker purple indicates a stronger predicted reversal of the disease signature. The list is enriched for symptom-modulating agents, including anti-inflammatories (fenoprofen) and hormonal therapies. **(B) Tahoe-100M Top Candidates.** Heatmap displaying the top 44 candidates identified using the single-cell Tahoe-100M database. In contrast to CMap, this list is heavily enriched for oncology-related kinase inhibitors and signaling modulators (e.g., pentoxifylline, futibatinib), reflecting the invasive, proliferative nature of endometriosis. **(A-B)** Black arrows **(>>> drug <<<)** highlight **irinotecan** and **terfenadine**, the only two drugs shared between the top rankings of both platforms. This minimal overlap illustrates the complementary nature of the resources: CMap captures established management strategies, while Tahoe-100M uncovers novel disease-modifying mechanisms akin to cancer therapy.

We subsequently evaluated the concordance between the two platforms by analyzing the top 50 drug candidates identified by each. The overlap was modest, highlighting the distinct yet complementary nature of the libraries. Among the top 50 hits in CMap, only six were present in Tahoe-100M, and conversely, only seven of the top 50 Tahoe hits were present in CMap. Despite these differences, we identified two drugs in common within the top rankings: terfenadine and irinotecan. This divergence suggests that the platforms capture different aspects of the disease biology. Tahoe-100M is heavily enriched for oncology and signaling inhibitors, surfacing numerous kinase inhibitors and epigenetic regulators such as FGFR inhibitors (pemigatinib, erdafitinib, futibatinib), CDK/AKT pathway drugs (dinaciclib, ipatasertib), and HDAC inhibitors (panobinostat, tucidinostat). This profile aligns well with modern characterizations of endometriosis as a proliferative, invasive, and inflammatory disease with cancer-like transcriptomic properties(*25, 26*). In contrast, CMap leans toward symptom modulation, prioritizing hormonal agents (levonorgestrel, medrysone), anti-inflammatory drugs (mesalazine, fenoprofen, resveratrol), and neuropsychiatric compounds (clomipramine).

The clinical relevance of these findings is supported by strong literature evidence for top hits from both pipelines. Both platforms successfully recovered established therapies: CMap identified levonorgestrel, a hormonal (progestin) therapy commonly used to treat endometriosis related pain, administered orally, as implants, and used in IUDs (*27–29*), while Tahoe-100M identified medroxyprogesterone acetate, which is widely used for hormonal suppression of lesions (Vercellini et al., 2003). Beyond standard of care, Tahoe-100M identified pentoxifylline as the top candidate common across all six disease signatures; its potential to prevent endometriosis recurrence after conservative surgery has been supported by controlled trials (*30, 31*). Similarly, CMap identified fenoprofen and resveratrol, both of which have documented efficacy in reducing lesion size and angiogenesis in animal models (*6, 32*). Several novel hits also show mechanistic promise; simvastatin (CMap) has been reported to inhibit stromal cell proliferation in vitro (*33*), while valproic acid (CMap) has shown potential in preclinical models to inhibit lesion growth (*25, 26, 34*). Furthermore, Tahoe-100M identified drospirenone, a progestin with anti-androgenic and anti-inflammatory properties relevant to pain control (*35*). While toxicity limits the clinical application of some hits, such as the topoisomerase inhibitor irinotecan (found in both pipelines), their identification underscores the pipeline’s ability to correctly identify antiproliferative mechanisms relevant to the disease pathology.

## DISCUSSION

### Principal Findings

The principal finding of this study is the high complementarity between CMap and Tahoe-100M: only 3.5% of recovered drugs were identified by both platforms, yet each captured a distinct and clinically relevant segment of the therapeutic landscape. While Tahoe-100M uniquely recovered 64.7% of candidates, capturing emerging targeted therapies like JAK and BTK inhibitors, CMap contributed 31.8% of hits, excelling at established therapeutics such as NSAIDs and corticosteroids. When benchmarked against Open Targets annotations, Tahoe-100M recovered a greater proportion of documented drug-disease associations than CMap (47.3% vs. 18.5%; 2.6-fold difference), with pronounced differences in autoimmune (4.3-fold) and oncology (3.0-fold) categories. These differences in annotated recovery rates should be interpreted cautiously, however: they partly reflect the greater overlap between Tahoe-100M’s drug library and Open Targets annotations, and the larger experimental breadth of Tahoe-100M (56,827 vs. 1,968 experiments capturing greater pharmacological and compound diversity), rather than representing a direct measure of superior predictive accuracy.

This divergence is driven by distinct mechanistic biases. Tahoe-100M displays a strongly “enzyme-centric” profile (44 - 46% of discoveries), reflecting its sensitivity to kinase cascades that create coherent transcriptomic signatures in oncology and inflammatory conditions. In contrast, CMap exhibits a “receptor-oriented” profile (35.8% membrane receptors), effectively capturing the rapid, membrane-proximal effects of agonists and ion channel modulators. These findings demonstrate that integrating heterogeneous perturbation resources is essential to maximize therapeutic coverage

Our disease category analysis provides actionable guidance for method selection. Tahoe-100M is recommended as the primary approach for oncology, autoimmune, and cardiovascular drug repurposing, where it demonstrated consistent 1.5 - 3.0-fold advantages. For metabolic and neurodegenerative diseases, CMap showed modest advantages, suggesting these categories may benefit from CMap-first or complementary approaches. The endometriosis case study illustrates an important caveat: for hormone-responsive conditions with established CMap-derived findings, CMap remains optimal for replication (62.5% recovery of Oskotsky et al.’s drugs), while Tahoe-100M provides value as a complementary discovery tool contributing unique candidates. This disease-specific performance pattern highlights that no single drug signature dataset is universally superior; method selection should be tailored to the disease biology and study objectives.

Our findings have several implications for computational drug repurposing practice, the first is that consensus predictions merit prioritization. Drugs identified by both CMap and Tahoe-100M showed 2.6-fold higher precision than single-method predictions, suggesting consensus hits represent higher-confidence candidates for experimental validation. Secondly, multi-method approaches maximize coverage. The high complementarity between methods (only 3.5–7.6% overlap in recovered drugs) indicates that single-method studies substantially underestimate the drug repurposing landscape. Integrating both CMap and Tahoe-100M captures distinct therapeutic signals. Lastly, platform-specific strengths inform therapeutic focus. CMap’s strength in established drug classes and Tahoe-100M’s identification of emerging targeted therapies suggest that platform selection can be tuned to study objectives, validation of known mechanisms versus discovery of novel interventions.

Our comparative analysis informs evidence-based platform selection for drug repurposing campaigns. Tahoe-100M is recommended for enzyme-driven pathologies (oncology, inflammation, metabolic disorders) and programs requiring high precision and biological concordance. CMap is optimal for receptor-mediated pathways (neurology, psychiatry, cardiovascular) and exploratory programs targeting ion channels or transporters. Given the minimal overlap (1.2%) yet distinct mechanistic strengths, we recommend an integrated strategy. Dual-platform hits represent high-confidence candidates suitable for prioritized validation, while platform-specific predictions should be pursued based on alignment between the disease mechanism and the platform’s established target-class bias.

Several limitations should be considered. First, validation relied on Open Targets, which creates a bias toward approved and late-stage candidates while missing off-label or early-stage therapeutics. Our definition of a true association, any drug evaluated for a given disease, favors sensitivity but may obscure specificity regarding efficacy. Second, despite standardized processing, disease signatures from CREEDS retain heterogeneity due to diverse experimental designs and tissue sources in the underlying GEO data. Third, our approach evaluates transcriptional reversal, which does not guarantee a specific therapeutic mechanism; observed reversals could stem from generalized stress responses rather than disease-specific correction. Fourth, many identified compounds, particularly from CMap, could not be mapped to Open Targets. This reflects the presence of proprietary or research-grade chemicals in CMap that lack public annotation, potentially leading to an underestimation of their relevance. Finally, the perturbation datasets differ technically: CMap uses bulk microarray profiling on limited cell lines, while Tahoe-100M utilizes single-cell RNA sequencing across a wider cellular panel. Although both rely heavily on neoplastic lines, they capture conserved biological processes relevant to non-cancer contexts. The superior resolution of the Tahoe-100M likely drives its improved precision. Ultimately, our findings suggest that CMap and Tahoe-100M are complementary, and platform selection should be driven by the specific translational context.

While this study identifies thousands of drug repurposing candidates validated against known therapeutics, bridging the gap to clinical application requires a rigorous translational framework (*36*). Top-ranked candidates, particularly those achieving “triple-validation” across CMap, Tahoe-100M, and prior literature, must first undergo experimental verification in high-fidelity models, such as patient-derived organoids or microphysiological systems, to confirm efficacy beyond standard cell lines (*37–39*). These efforts should be complemented by deep mechanistic profiling to ensure that signature reversal is driven by therapeutically relevant target engagement rather than off-target toxicity, a step particularly critical for drugs with polypharmacology (*40*). For candidates already approved for other indications, this translation can be accelerated by systematically reviewing real-world evidence and electronic health records to de-risk safety profiles in new patient populations (*41*). Furthermore, our observation that Tahoe-100M achieves variable recovery rates across autoimmune conditions underscores the critical need for transcriptomic patient stratification to identify responsive subtypes (*42*). Ultimately, randomized clinical trials remain the definitive gold standard; future studies must transition validated candidates into Phase II proof-of-concept or innovative basket trials to establish efficacy and safety in human subjects prior to broad clinical implementation (*43*).

This comprehensive evaluation demonstrates that Tahoe-100M and CMap provide complementary perspectives on the drug repurposing landscape, with Tahoe-100M showing overall superior recovery of known therapeutics and particular strength in oncology and autoimmune conditions. The high complementarity between methods, with only 3.5–7.6% overlap in recovered drugs, argues for multi-method approaches as standard practice in computational drug repurposing. Consensus predictions from both platforms represent high-confidence candidates meriting experimental prioritization. Our disease category-level recommendations and case study analyses provide actionable guidance for method selection across diverse therapeutic areas.

## MATERIALS AND METHODS

### Disease and Drug Signatures

This study was designed to systematically compare and integrate two connectivity mapping approaches for transcriptional drug repurposing. We conducted a retrospective computational analysis using publicly available disease signatures and drug perturbation profiles, with validation against known drug-disease associations. The primary outcomes were recall (recovery rate) of known therapeutics and identification of consensus candidates across platforms. Sample sizes were determined by the availability of curated disease signatures (n=233) in CREEDS and their matchability to Open Targets (n=203 evaluable diseases).

Disease signatures were obtained from CREEDS, a manually curated database of 233 disease-associated differential expression signatures derived from Gene Expression Omnibus (GEO) studies (***Fig. 8***) (*18*). Each signature contains up-regulated and down-regulated gene sets representing transcriptomic changes associated with disease states relative to healthy controls. Signatures were quality-controlled during original curation; we applied no additional filtering prior to analysis.

We used the original Connectivity Map (CMap) resource containing bulk microarray perturbation profiles. Following quality control filtering based on internal consistency (Pearson r ≥ 0.15 between replicate signatures), 1,968 of 6,100 original experiments (32.3%) were retained for analysis. The filtered dataset covered 13,071 genes across multiple cell lines and drug concentrations.

We used pseudo-bulk perturbation profiles derived from large-scale single-cell RNA sequencing experiments (Tahoe-100M). The Tahoe-100M resource contained 56,827 experiments profiling drug responses across diverse cell types and conditions. Gene-level significance filtering (p ≤ 0.05) was applied during signature generation to retain only significantly perturbed genes. For connectivity scoring, the retained log2 fold-change values were converted to genome-wide ranks: all genes in each experiment’s signature were ranked by their log2 fold-change, and these ranks served as the input for the nonparametric KS-based scoring procedure (see below).No additional experiment-level filtering was required due to naturally high data quality (***Fig. 1***). After mapping to a shared gene universe, 22,168 genes were available for analysis.

### CDRPipe Computational Pipeline

Disease signatures were standardized by mapping gene symbols to Entrez IDs using g:Profiler (gconvert function, organism = “hsapiens”, target = “ENTREZGENE_ACC”). Genes were filtered by adjusted p-value (< 0.05, when available) and absolute log fold-change (> 0.25). Only genes present in the respective drug perturbation database gene universe were retained.

For each disease signature and drug perturbation profile, we computed a modified Kolmogorov-Smirnov (KS) connectivity score quantifying transcriptional reversal. Briefly, disease up-regulated genes were expected to be ranked low in drug signatures (indicating downregulation by drug), while disease down-regulated genes were expected to be ranked high (***Supplement Fig. 1***). The connectivity score combines KS statistics for up and down gene sets:

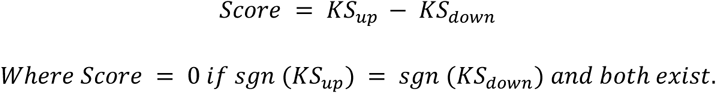

Negative scores indicate transcriptional reversal (drug reverses disease signature); positive scores indicate mimicry.

Statistical significance was assessed using an empirical null distribution generated from 100,000 random gene sets matched in size to the query disease signature. Two-sided p-values were computed as the proportion of random scores with absolute value exceeding the observed score. To prevent zero p-values, we applied the permutation-based minimum: p = 1/(N+1) when no random scores exceeded the observed value (Phipson & Smyth, 2010). False discovery rate correction was performed using the qvalue package; when qvalue failed due to distribution assumptions, Benjamini-Hochberg adjustment was applied.

Drug candidates were identified using a significance threshold of q < 0.05 combined with negative connectivity score (reversal direction). For each disease, the most significant instance per drug (minimum score across cell lines and concentrations) was retained to avoid duplicate counting.

### Validation Against Known Drug-Disease Associations

To validate our pipeline against established clinical knowledge, we obtained known drug-disease associations from the Open Targets Platform (version 2024). Disease entities from CREEDS were systematically mapped to Open Targets using a hierarchical approach, identifying 151 diseases (64.8%) through exact name matching and 52 diseases (22.3%) via ontology synonyms, while 30 diseases (12.9%) remained unmatched (***Table 2***). Performance was subsequently evaluated by calculating Recall (*S*/*P x* 100), defined as the proportion of available known drugs in the perturbation database (*P*) that were successfully recovered by CDRPipe (*S*) with a significant reversal score (*q* < 0.05).

**Table 2.** Classification of Disease Signatures by Therapeutic Area. Classification of disease signatures into 20 therapeutic area categories based on Open Targets disease ontology annotations. Of the 233 disease signatures obtained from CREEDS, 203 (87.1%) were successfully matched to Open Targets through exact name or synonym matching, and 180 of these had at least one known drug association available for validation. Categories are ranked by disease count within the final analysis cohort (n=180). Each disease was assigned to a single primary therapeutic area. Representative examples are provided for each category.

| Therapeutic Area | N | % | Examples |
| --- | --- | --- | --- |
| Cancer/Tumor | 27 | 15.0% | Lung adenocarcinoma, Colorectal cancer, Melanoma, Glioblastoma multiforme, Ovarian cancer |
| Genetic/Congenital | 20 | 11.1% | Cystic fibrosis, Duchenne muscular dystrophy, Down syndrome, Huntington's disease, Marfan syndrome |
| Nervous System | 18 | 10.0% | Alzheimer's disease, Multiple sclerosis, Epilepsy syndrome, Amyotrophic lateral sclerosis,... |
| Immune System | 13 | 7.2% | Rheumatoid arthritis, Psoriasis, Sjögren's syndrome, Ankylosing spondylitis, Psoriatic arthritis |
| Gastrointestinal | 11 | 6.1% | Ulcerative colitis, Crohn's disease, Barrett's esophagus, NASH, Gastroesophageal reflux disease |
| Musculoskeletal | 11 | 6.1% | Osteoarthritis, Systemic lupus erythematosus, Scleroderma, Osteoporosis, Arthritis |
| Cardiovascular | 9 | 5.0% | Myocardial infarction, Hypertension, Atherosclerosis, Dilated cardiomyopathy, Pulmonary hypertension |
| Infectious Disease | 9 | 5.0% | Tuberculosis, Influenza, Hepatitis C, HIV infectious disease, Dengue disease |
| Respiratory | 8 | 4.4% | Asthma, Chronic obstructive pulmonary disease, Allergic asthma, Acute lung injury, Bronchopulmonary dysplasia |
| Hematologic | 7 | 3.9% | Sickle cell anemia, Aplastic anemia, Myelodysplastic syndrome, Chronic myeloid leukemia, Acute T cell leukemia |
| Psychiatric | 7 | 3.9% | Schizophrenia, Autism spectrum disorder, Alcohol abuse, Nicotine dependence, Bipolar disorder |
| Reproductive/Breast | 7 | 3.9% | Breast cancer, Endometriosis, Prostate cancer, Endometrial cancer, Polycystic ovary syndrome |
| Endocrine System | 6 | 3.3% | Polycystic ovary syndrome, Papillary thyroid carcinoma, Adrenoleukodystrophy, Androgen insensitivity syndrome |
| Phenotype | 6 | 3.3% | Ischemic stroke, Septic shock, Hypoxia, Dystonia, Intellectual disability |
| Urinary System | 5 | 2.8% | Renal cell carcinoma, Cystitis, Nephroblastoma, Interstitial cystitis, Diabetic nephropathy |
| Skin/Integumentary | 5 | 2.8% | Atopic dermatitis, Eczema, Psoriasis vulgaris, Actinic keratosis, Urticaria |
| Metabolic | 5 | 2.8% | Diabetes mellitus, Obesity, Morbid obesity, MELAS syndrome, Familial hypercholesterolemia |
| Pancreas | 3 | 1.7% | Type 1 diabetes mellitus, Type 2 diabetes mellitus, Pancreatic ductal adenocarcinoma |
| Visual System | 2 | 1.1% | Glaucoma, Primary open angle glaucoma |
| Pregnancy/Perinatal | 1 | 0.6% | Pre-eclampsia |

### Case Study Analyses

For the autoimmune disease case study, 18 autoimmune diseases with available Open Targets matches were analyzed in depth. For each disease, CDRPipe was run independently against both CMap and Tahoe-100M perturbation libraries using the standard pipeline parameters (q < 0.05, negative connectivity score). Drug-level analysis classified each recovered drug by its platform source (CMap only, Tahoe-100M only, or both). Clinical validation was performed by cross-referencing recovered drugs against Phase 4 clinical trial data from Open Targets for each specific disease indication. Recovery rates were compared between platforms using the Wilcoxon signed-rank test, with effect size quantified by Cohen’s d.

For the endometriosis case study, six disease signatures from Oskotsky et al. (Unstratified, Stage I-II, Stage III-IV, Proliferative Phase, Early Secretory Phase, Mid-Secretory Phase) were re-analyzed using both CMap and Tahoe-100M. To ensure exact replication of the original findings, the CMap analysis used the parameters reported in Oskotsky et al.: p-value < 0.05, absolute log2 fold change > 1.1, random seed = 2009, and 1,000 permutations. The same parameters were then applied to Tahoe-100M for direct comparison. Top candidates were ranked by connectivity score across signatures, and concordance between platforms was assessed by comparing the top 50 candidates from each.

Diseases were categorized into nine therapeutic areas using keyword matching: Oncology, Metabolic, Neurodegenerative, Cardiovascular, Infectious, Autoimmune, Allergic/Respiratory, Rare/Genetic, and Other. Category-level performance was summarized as mean known drug hits per disease.

### Statistical Analysis

All statistical analyses were performed in R (version 4.2+). Paired comparisons of Tahoe-100M versus CMap recovery rates were performed using the Wilcoxon signed-rank test. Effect sizes were quantified using Cohen’s d. P-values < 0.05 were considered statistically significant. For the overall recovery rate comparison, 153 diseases with evaluable data for both platforms were included in the paired analysis. Complementarity was quantified as the percentage of recovered drugs identified by both platforms, Category-level recommendations were based on mean known drug hits with a threshold of >2 drugs difference for primary method recommendation.

## Funding

National Institutes of Health grant 1R21HD114953 (MS, LCG)

National Institutes of Health grant 1R01AI180118 (MS)

National Institutes of Health grant P30 AR070155 (MS)

National Institutes of Health grant P01HD106414 (LCG)

National Institutes of Health grant 1R01AG100879-0 (MS)

UCSF March of Dimes Prematurity Research Center

## Author contributions

Conceptualization: EN, UK, MS

Methodology: EN, UK, MS, JN, LCG, TO

Investigation: EN

Software: EN, BO

Data curation: EN, UK, XT, LA, EC, BA, CPS

Formal analysis: EN, UK, XT, LA, EC, BA, CPS

Visualization: EN

Supervision: MS

Funding acquisition: MS, LCG

Writing – original draft: EN, MS

Writing – review & editing: UK, XT, LA, EC, BA, CPS, BO, BG, DKS, JN, LCG, TO

## DATA and MATERIALS AVAILABILITY

- The CDRPipe R package and R Shiny app is available: https://github.com/enockniyonkuru/drug_repurposing
- The CDRPipe Comparative Analysis for this manuscript: https://github.com/enockniyonkuru/cdrpipe-comparative-analysis
- The R Shiny app is freely accessible via https://cdrpipe.org/.
- Disease signatures from CREEDS are publicly available: https://maayanlab.cloud/CREEDS/
- Open Targets drug-disease associations are available: https://platform.opentargets.org/
- CMap perturbation profiles are available from the Broad Institute: https://www.broadinstitute.org/connectivity-map-cmap
- Tahoe-100M perturbation profiles are available from hugging face: https://huggingface.co/datasets/Tahoe-100Mbio/Tahoe-100M-100M

## SUPPLEMENT

**Supplement Algorithm 1:**
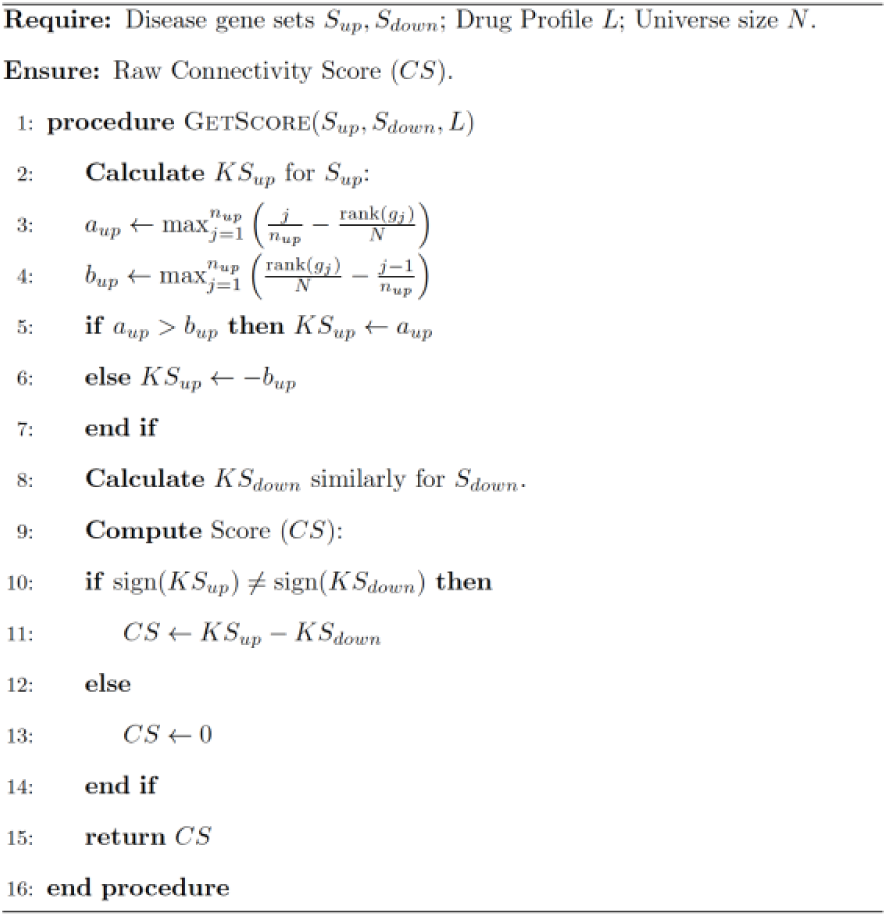
Calculation of Connectivity Score.

**Supplement Algorithm 2:**
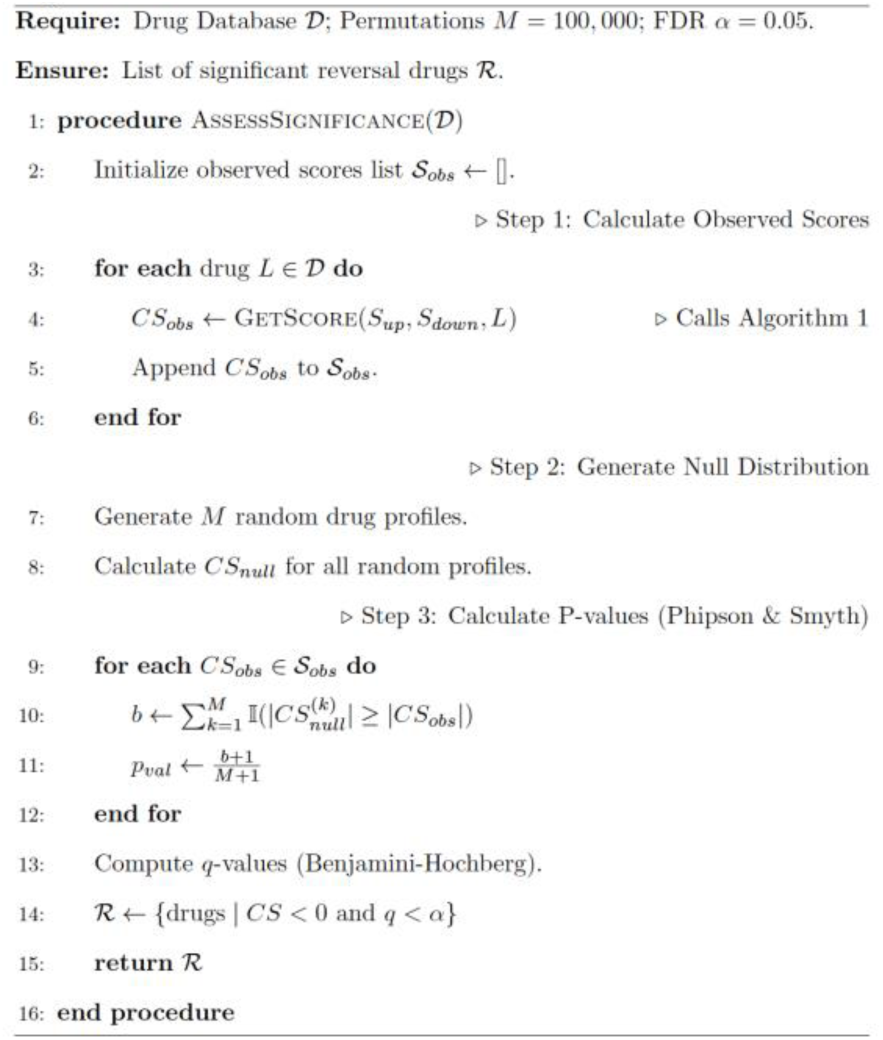
Statistical Assessment and Candidate Selection.

**Supplement Fig. 1.**
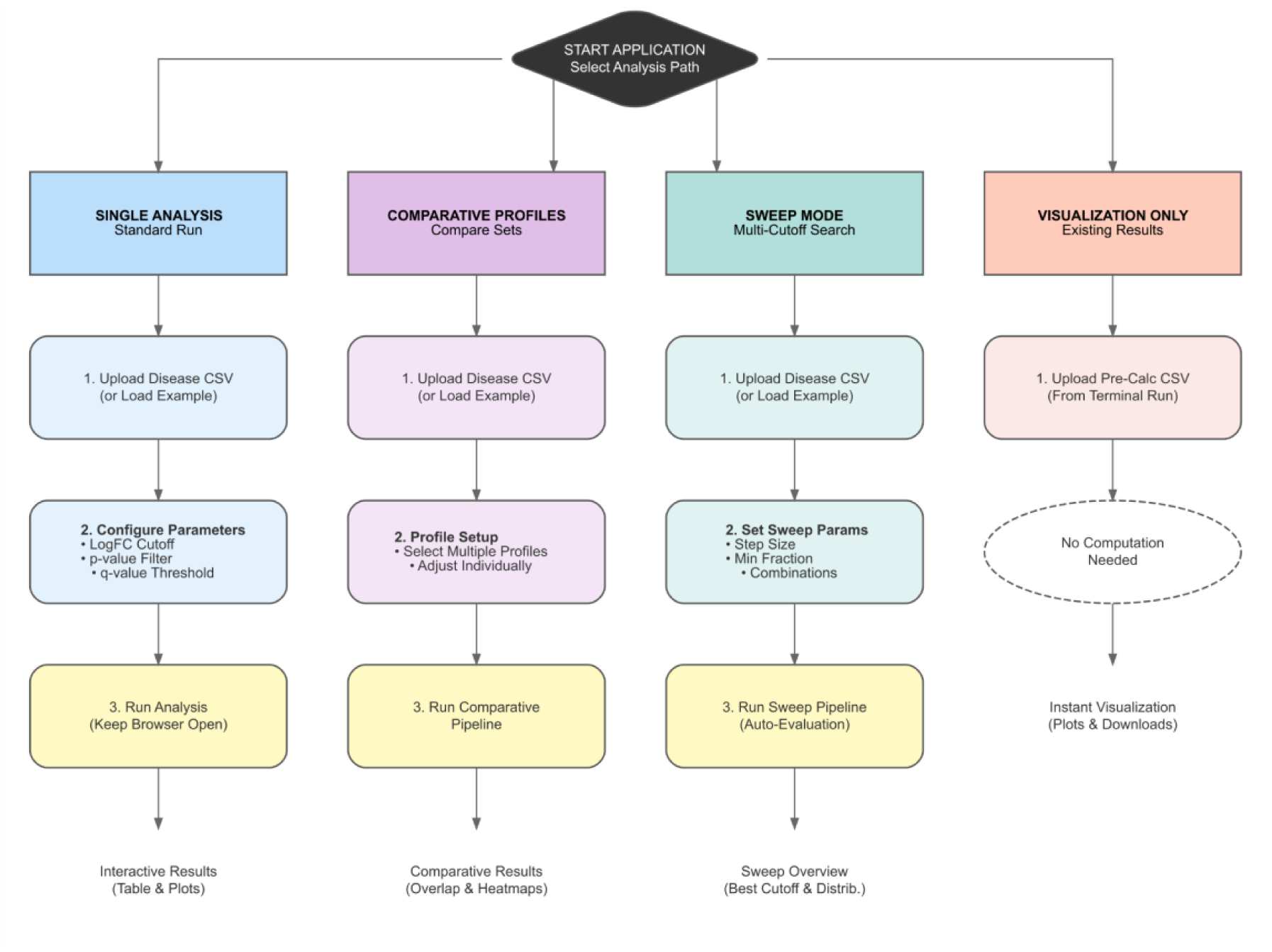
CDRPipe Shiny Application Workflows. The application interface guides users through four distinct analysis modules starting from the main dashboard: (**1**) **Single Analysis:** A standard workflow for running drug repurposing on individual disease signatures. (**2**) **Comparative Profiles:** A module for analyzing and contrasting multiple disease sets simultaneously to identify shared or distinct therapeutic signals. (**3**) **Sweep Mode:** An optimization tool that performs a multi-cutoff grid search to identify ideal parameter thresholds. (**4**) **Visualization Only:** A rapid-access mode for rendering plots from pre-computed results files without re-running the computational pipeline. Each path follows a structured workflow of data upload, parameter configuration, execution, and interactive result exploration.

**Supplement Fig. 2.**
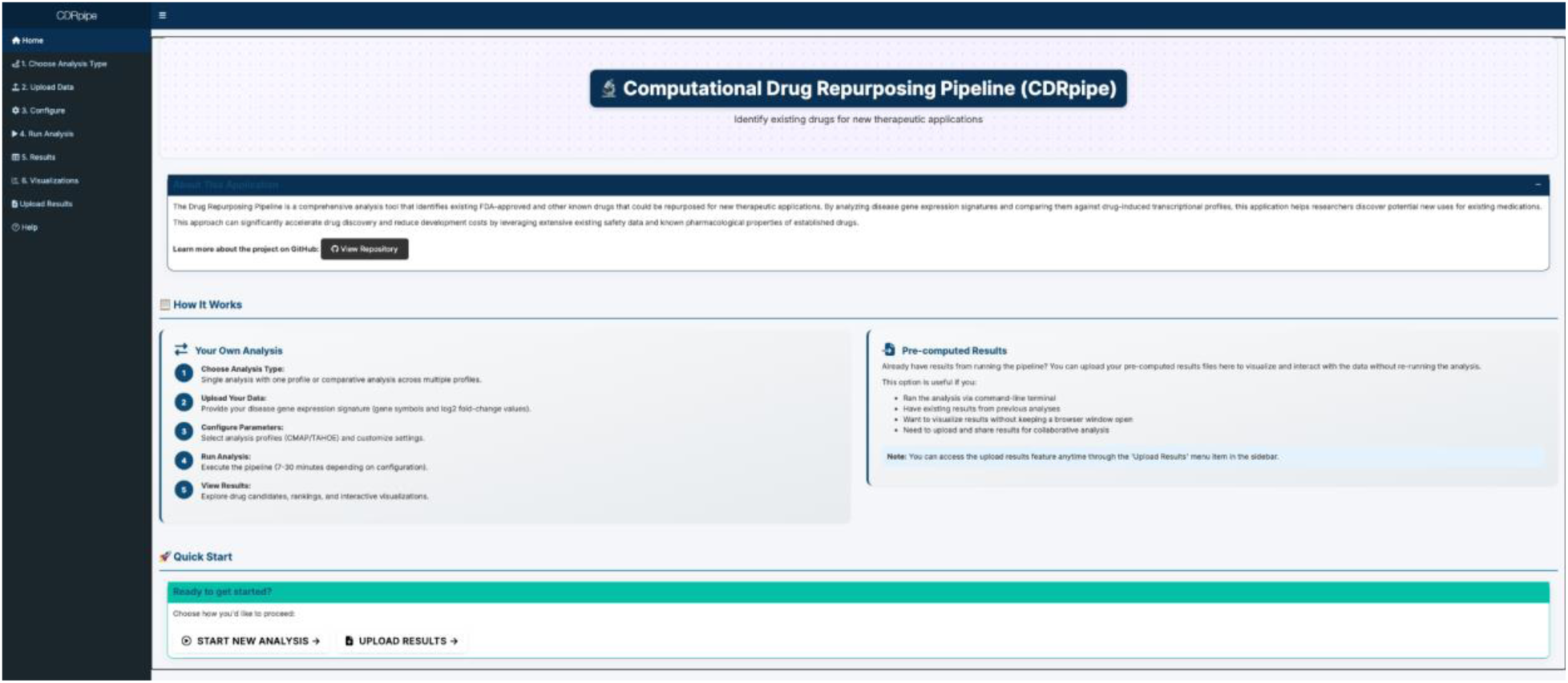
CDRPipe R Shiny Application Home Page. The figure displays the landing interface of the web application, which serves as the central hub for the DRPipe workflow. The dashboard features a **“How It Works”** section that outlines the step-by-step process for conducting a new analysis, from choosing an analysis mode and uploading gene expression signatures to configuring parameters and interpreting results. Additionally, the interface offers a **“Pre-computed Results”** pathway, allowing users to upload and visualize existing pipeline outputs without re-executing the computational steps. Navigation is facilitated through the sidebar menu and **“Quick Start”** action buttons.

**Supplement Fig 3:**
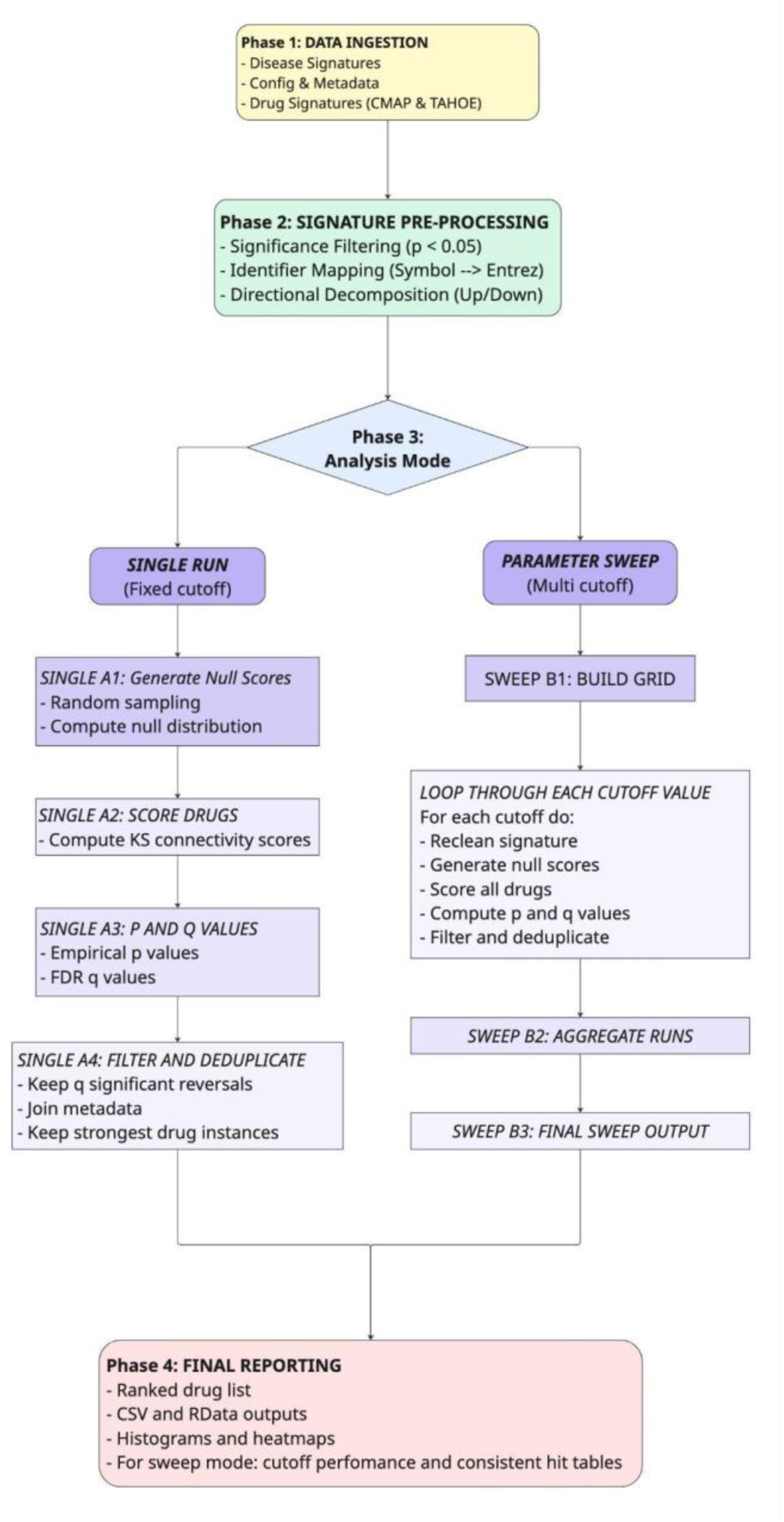
Schematic overview of the CDRPipe workflow. The pipeline proceeds in four distinct phases: (**1**) **Data Ingestion** of disease signatures and drug perturbation profiles (CMAP and Tahoe-100M). (**2**) **Signature Pre-processing**, which standardizes inputs via significance filtering (p < 0.05), identifier mapping to Entrez IDs, and directional decomposition into up/down gene sets. (**3**) **Analysis Mode**, where users select between a **Single Run** (using fixed cutoffs) or a **Parameter Sweep** (optimizing thresholds via grid search). This phase executes the core algorithm: generating null distributions, computing Kolmogorov-Smirnov (KS) connectivity scores, and assessing statistical significance via empirical p-values and FDR correction. **(4) Final Reporting**, which outputs ranked candidate lists, visualization plots, and comprehensive result tables for downstream validation.

**Supplement Table 1:** Autoimmune Disease Summary. **Comparative Performance Metrics Across 18 Autoimmune Diseases.** The table provides a detailed breakdown of drug recovery statistics for each disease analyzed in Case Study 1. Columns quantify the Known Drugs (total validated treatments in Open Targets), the subset Available within each platform’s library, the total number of significant predictions (Hits), and the number of validated predictions (Recovered) for both CMAP and Tahoe-100M. The Recovery Rate columns highlight the performance disparity between the platforms; notably, Tahoe-100M achieves 100% recovery of available known drugs for 8 of the 18 conditions (e.g., Psoriatic Arthritis, Inclusion Body Myositis), whereas CMAP frequently shows 0% recovery for the same indications.

| Disease | Known Drugs (DB) | Available (CMAP) | Hits (CMAP) | Recovered (CMAP) | Recovery Rate (CMAP) | Available (TAH OE) | Hits (TAH OE) | Recovered (TAH OE) | Recovery Rate (TAH OE) | Total Candidates | Total Recovered | Overall Recovery Rate |
| --- | --- | --- | --- | --- | --- | --- | --- | --- | --- | --- | --- | --- |
| Scleroderma | 18 | 4 | 300 | 2 | 50.00% | 1 | 230 | 1 | 100.00% | 978 | 3 | 16.67% |
| Autoimmune thrombocytopenic purpura | 53 | 16 | 28 | 2 | 12.50% | 11 | 379 | 11 | 100.00% | 474 | 13 | 24.53% |
| Psoriasis | 149 | 20 | 273 | 15 | 75.00% | 14 | 332 | 14 | 100.00% | 574 | 25 | 16.78% |
| Psoriatic arthritis | 47 | 9 | 5 | 0 | 0.00% | 6 | 365 | 6 | 100.00% | 709 | 6 | 12.77% |
| Inclusion body myositis | 8 | 1 | 7 | 0 | 0.00% | 1 | 357 | 1 | 100.00% | 419 | 1 | 12.50% |
| Discoid lupus erythematosus | 9 | 1 | 6 | 0 | 0.00% | 2 | 357 | 2 | 100.00% | 433 | 2 | 22.22% |
| Dermatomyositis | 29 | 4 | 93 | 0 | 0.00% | 2 | 370 | 2 | 100.00% | 463 | 2 | 6.90% |
| Sjogren's syndrome | 39 | 6 | 701 | 4 | 66.67% | 4 | 315 | 4 | 100.00% | 1086 | 8 | 20.51% |
| Inflammatory bowel disease | 33 | 11 | 109 | 1 | 9.09% | 7 | 345 | 6 | 85.71% | 503 | 6 | 18.18% |
| Ankylosing spondylitis | 43 | 12 | 7 | 0 | 0.00% | 9 | 272 | 7 | 77.78% | 417 | 7 | 16.28% |
| Crohn's disease | 104 | 27 | 102 | 2 | 7.41% | 12 | 332 | 9 | 75.00% | 476 | 11 | 10.58% |
| Relapsing-remitting multiple sclerosis | 79 | 19 | 679 | 6 | 31.58% | 9 | 248 | 6 | 66.67% | 1065 | 12 | 15.19% |
| Rheumatoid arthritis | 240 | 45 | 164 | 8 | 17.78% | 25 | 272 | 16 | 64.00% | 859 | 22 | 9.17% |
| Systemic lupus erythematosus | 103 | 20 | 268 | 8 | 40.00% | 15 | 267 | 9 | 60.00% | 894 | 16 | 15.53% |
| Ulcerative colitis | 115 | 24 | 68 | 3 | 12.50% | 10 | 207 | 5 | 50.00% | 619 | 8 | 6.96% |
| Multiple sclerosis | 145 | 43 | 247 | 12 | 27.91% | 16 | 141 | 7 | 43.75% | 774 | 18 | 12.41% |
| Juvenile idiopathic arthritis (sJIA) | 10 | 11 | 72 | 0 | 0.00% | 8 | 81 | 3 | 37.50% | 191 | 3 | 30.00% |
| Type 1 diabetes mellitus | 138 | 31 | 201 | 8 | 25.81% | 12 | 178 | 2 | 16.67% | 807 | 10 | 7.25% |

**Supplement Table 2: CMap Drugs List**: https://drive.google.com/file/d/1-_eaCW600Tu7tE-cDOyqqC1Xn8KUprof/view?usp=sharing

**Supplement Table 3: Tahoe-100M Drugs List:** https://drive.google.com/file/d/1TsjJeYhW-5p_BiENejMCv2EWiSPww0Fd/view?usp=sharing

**Supplement Table 4: Tahoe-100M & CMap Drug Overlap List:** https://drive.google.com/file/d/13lZhEOnQF0sCBLTutsASkUW-C8lg9Z_T/view?usp=sharing

**Supplement Table 4: CREEDs Disease Names List:** https://drive.google.com/file/d/1-hL2SkOfaylUiotNCU73zlv9bkxMAq0q/view?usp=sharing

